# CluSem: Accurate Clustering-based Ensemble Method to Predict Motor Imagery Tasks from Multi-channel EEG Data

**DOI:** 10.1101/2021.09.05.458710

**Authors:** Md. Ochiuddin Miah, Rafsanjani Muhammod, Khondaker Abdullah Al Mamun, Dewan Md. Farid, Shiu Kumar, Alok Sharma, Abdollah Dehzangi

## Abstract

**Background:** The classification of **m**otor **i**magery **e**l**e**ctroencephalo**g**ram (MI-EEG) is a pivotal task in the biosignal classification process in the **b**rain-**c**omputer **i**nterface (BCI) applications. Currently, this bio-engineering-based technology is being employed by researchers in various fields to develop cutting-edge applications. The classification of real-time MI-EEG signals is the most challenging task in these applications. The prediction performance of the existing classification methods is still limited due to the high dimensionality and dynamic behaviors of the real-time EEG data.

**Proposed Method:** To enhance the classification performance of real-time BCI applications, this paper presents a new clustering-based ensemble technique called CluSem to mitigate this problem. We also develop a new brain game called CluGame using this method to evaluate the classification performance of real-time motor imagery movements. In this game, real-time EEG signal classification and prediction tabulation through animated balls are controlled via threads. By playing this game, users can control the movements of the balls via the brain signals of motor imagery movements without using any traditional input devices.

**Results:** Our results demonstrate that CluSem is able to improve the classification accuracy between 5% and 15% compared to the existing methods on our collected as well as the publicly available EEG datasets. The source codes used to implement CluSem and CluGame are publicly available at https://github.com/MdOchiuddinMiah/MI-BCI_ML.

## 1. Introduction

Brain engineering is an emerging field in the science and technology. This multidisciplinary branch consists of the combination of various fields such as physics, chemistry, biology, computer science, and mathematics [1]. It also incorporates the concepts and methods of other fields such as clinical medicine and engineering [2]. Brain-Computer Interface (BCI) is a branch in brain engineering that aims to solve practical problems of the life sciences. BCI and Human Machine Interface (HMI) are the modern technologies that are used as tools to establish communication between users and machines [3].

BCI technology incorporates neurophysiological activities as input signals and interprets them into meaningful commands employing Machine Learning (ML) algorithms [4]. BCI technology has brought tremendous opportunities in the medical field especially for the treatment and rehabilitation of people with disabilities [5, 6]. Around the world, millions of people are affected by various forms of disabilities and physical impairments due to different reasons such as childhood disability, old-age disability, early-age disability, accidental/unforeseen issues, and serious health problems [7]. For people with disabilities, it is extremely difficult or, in some cases, impossible to do day-to-day chores without any assistance from a caregiver.

A wide variety of equipment and devices are invented in this field for rehabilitating and aiding people with disabilities [8, 5]. This technology allows people with disabilities to command and control external devices such as computers, wheelchairs, and robots by utilizing their thoughts. Notwithstanding, the horizon of this field has been broadened to a wide range of non-medical applications such as gaming, entertainment, military, and meditation training [9, 10]. The knowledge of BCI can also be applied in smart environment systems like smart houses or smart workplaces. This technology has also opened a new window of opportunities for the gaming and entertainment industry. Furthermore, different types of games already being developed using this technology including games to relieve users from stress [11, 12].

Fundamentally, BCI technology can be employed in three ways based on the process of signal acquisition from brain [13]. First is to place wires inside the grey matter of the brain to intensify brain signals. This method is called invasive BCI [14, 3]. The second way is to place electrodes on the surface of the scalp to measure activities from a huge group of neurons. This method is called non-invasive BCI [15, 16]. In comparison with invasive BCI, this method is less sophisticated but at the same time, it does not intensify the input signals as good [17, 18]. The third type of BCI is called partially invasive BCI. In this technique, wires are placed inside the brain but above the grey matter [13, 17].

During the past few years, a wide range of machine learning methods including ensemble classifiers has been used for motor imagery EEG signal classification. We investigated assessed the performance of existing single and ensemble classifiers in real-time BCI applications in our prior studies [17, 14, 19]. It is well-known that the existing classifiers are not very accurate due to the high dimensionality and dynamic behaviors of the real-time EEG data [20, 21]. The signals may be biased with artifacts and noise due to the low conductivity of the electrodes with the scalp [1, 6].

The objective of this paper is to extend our prior works and ameliorates the classification performance by handling multiple electrodes or neurons data at the same time and proposing a cluster-based ensemble classifier. Our proposed model, which is called CluSem, first clusters the input data based on the position of the electrodes. It also selects the model dynamically based on the electrode locations to classify real-time EEG data. It then uses an ensemble classifier to classify the motor imagery EEG signals. We compare CluSem with different classifiers that have been widely used to tackle this problem such as Artificial Neural Network (ANN), Support Vector Machine (SVM), Naïve Bayes, Decision Tree, Random Forest (RF), Bagging, and AdaBoost. Our results illustrate that CluSem can significantly outperform those models. CluSem demonstrates between 5% and 15% prediction enhancement over those existing methods found in the literature.

Besides, we also developed a new brain game called CluGame to distinguish human thoughts in real-time based on CluSem. To develop this game, we employed the threading technique to control the signal classification and prediction tabulation via animated ball in real-time. The CluGame can be used for rehabilitation as well as upgrading user well-being. The user can exercise its concentration to recover from attention deficiency and boost attention via playing this brain game. CluGame can also be used for gaming and entertainment purposes. The source codes of the CluSem and the CluGame are publicly available at https://github.com/MdOchiuddinMiah/MI-BCI_ML/tree/master/Brain-Game. We also generate new MI-EEG brain data for our experiments which is also publicly available at https://github.com/MdOchiuddinMiah/MI-BCI_ML/tree/master/Datasets.

The remainder of this paper is organized as follows: Section 2 presents the related works. Section 3 illustrates brain measurement techniques, signal acquisitions, pre-processing, feature extraction, and datasets descriptions. Section 4 covers supervised classification and the proposed clustering-based ensemble method CluSem. Section 5 provides experimental results and introduces CluGame. Finally, Section 6 presents conclusions and future direction.

## 2. Related Works

During the last decade, a wide range of machine learning approaches has been proposed to classify the motor imagery EEG signal [17, 20, 20, 22]. In real-time applications, the reported prediction performance has remained relatively low due to the high dimensionality and dynamic behaviors of the real-time EEG data [4, 23, 21, 24]. In the MI EEG signal classification study, Kumar et al. [25] employed a mutual information-based frequency band selection approach to utilize all information that is got from different channels. Furthermore, they introduced a sub-band between 7 and 30 Hz to cover a wide range of frequencies. To extract features, linear discriminant analysis is used for each filter bank. Extracted motor imagery EEG data are classified by a popular algorithm named support vector machine. In another study, Kumar et al. [26] proposed a common spatial-spectral pattern (CSSP) and optimization of temporal filters to enhance MI EEG classification accuracy. To augment the spatial resolution, CSSP is introduced. On the other hand, a temporal filter is used to optimize every subject for each frequency band that carries informative information. Later on, Kumar et al. [27] proposed a spatial-frequency-temporal feature extraction (SPECTRA) tool that will find features from real-time BCI systems. It shows enhancement in brain wave signal recognition and outperformed other competing methods using three public benchmark datasets.

Recently, Raza et al. [1] presented an adaptive ensemble approach to EEG classification to handle non-stationarity in the motor imagery task. Their proposed model collected MI correlated brain responses and extracted spatial pattern structures. They then developed their ensemble classifier, to overcome the deviations in streaming data input. Their results demonstrated that using an Ensemble classifier obtains better results than using single classifiers. In the same year, Sreeja et al. [28] proposed a sparse based classification method to classify MI related tasks from EEG data. To reduce the computational time, they only used wavelet energy without any pre-processing as a feature to classify MI data. They showed that their proposed approach outperforms existing classifiers.

Later on, Li et al. [2] proposed a novel decoding approach employing Overlapping Averaging (OA) to interpret MI-EEG data. To overwhelm the constraint of the general Region of Interest (ROI) based decrypting approach, they used Weighted Minimum Norm Estimate (WMNE). They studied their model on a public dataset and achieved higher decrypting accuracy than existing approaches. In a different study, Pattnaik and Sarraf [29] attempted to apply left and right-hand movement classification using raw EEG signals. Before applying the movement classification, they removed the artifacts in the obtained signals employing a low pass filtering technique.

Recently, Shanechi [22] examined the decoding algorithms made in the BMI study. They hypothesized that it is possible to design a motor BMI as a closed-loop control system. They inspected current decoder designs that emphasize the unique properties of BMI. Moreover, they discussed possible opportunities to formulate a control-theoretic framework to design BMI, as well as, the development of more advanced BMI control algorithms. In the same year, Mehmood et al. [30] proposed a method that considerably enhances the rate of emotion recognition for the popularly implemented spectral power band routine. Features selected by this routine performed better than both univariate and multivariate attributes. The optimal attributes were later processed to classify emotion by applying KNN, SVM, LDA, Naïve Bayes, RF, and Deep Learning (DL) classifiers.

In a different study, Mamun et al. [19] introduced a new technique to procure frequency reliant neural synchronization as well as inter-hemispheric connectivity properties. They based their method on Granger causality as well as Wavelet Packet Transform (WPT) approaches. Their approach is capable of decoding movement associated behaviors accurately, from the registered Local Field Potentials (LFPs) activity. The finding of this study shows that nominated optimum neural synchronization associated with inter-hemispheric connectivity has the potential to capture signals to supplement the adaptive Brain-Machine Interface (BMI). Furthermore, Lu et al. [31] investigated the prospect of ameliorating performance in a Transcranial Doppler ultrasonography (TCD) based BCI. They used the structures and classifiers that are computationally suitable for online application by running an offline investigation of TCD recordings and obtained promising results. Combining SVM classifiers with Weighted Sequential Forward Selection (WSFS), they outperformed similar studies found in the literature.

Afterward, Sun et al. [20] investigated a wide range of ensemble classifiers such as bagging, boosting, and random subspace to classify MI related tasks from EEG signals. They proposed an ensemble of three classifiers namely, SVM, DT, and K-Nearest Neighbors (k-NN) to solve this problem and achieved promising results. Later on, Lebedev and Nicolelis [32] reviewed some of the essential challenges encountered by BMI research. They proposed a chain of milestones to convert up to date experimental advances into viable medical applications within the coming 10 to 20 years. The guideline they provided underscores the contemporary history of the BMI and puts a strong emphasis on the influential factors related to its growth.

## 3. EEG Signal Acquisition

### 3.1. Functional Areas of Brain

The human brain is the main organ in the nervous system [19]. The central nervous system has consisted of both the brain and spinal cord. The human brain acts as the master of the whole body, as it controls almost all of the body activities. It performs different functions such as receiving, processing, and generating information [23]. It acts as a receiver for the sensory organs sending information to the brain. The brain then integrates, coordinates, and processes the given input information to produce decisions and instructions to the rest of the body [29].

Different brain parts are responsible for completing different tasks. Among them, the motor system is the part that both generates and controls the movements of the body [19]. Also, the nerves do the job of transferring the motor system-generated movements from the brain to the motor neurons in the body. In fact, the action of the muscle is governed by this process. Using the spinal cord, the corticospinal tract passes movements to the torso as well as to the limbs. The eyes, mouth, and face related movements are carried by the cranial nerve [32].

The motor cortex generates the movement of arms and legs. It consists of three parts namely, primary motor cortex, premotor cortex, and supplementary motor area [31]. Locating on the frontal lobe of the brain, the primary motor cortex is one of the essential areas that are necessary for motor function. Additionally, the primary motor cortex produces neural impulses and then these impulses control the execution of movement [14].

### 3.2. Brain Measurement Techniques

In BCI, different kinds of neurological modalities are applied to obtain neurological brain signals [19]. Electroencephalography (EEG) is one of the popular methods for measuring brain activities. It is commonly a non-invasive approach and assesses voltage oscillations ensuing from ionic current inside the brain neurons [13]. Other non-invasive methods for measuring brain activities are positron emission tomography (PET), magneto-encephalography (MEG), Transcranial Doppler ultrasonography (TCD), and functional magnetic resonance imaging (fMRI) [10, 33]. Some of the invasive electrophysiological methods are namely, local field potentials (LFPs), electrocorticography (ECoG), and single-unit recording [19, 14]. EEG, ECoG, LFPs, and single-neuron recordings are popular because they are relatively simple and less costly. They also produce, high temporal resolution [19, 34]. On the other hand, fMRI, MEG, and PET are very costly and time-consuming [13]. In our experiment, we used Emotiv EPOC+ EEG neuroheadset because it is less sophisticated, more cost-efficient, and easy to use [17, 14].

### 3.3. EEG Emotiv EPOC+ Neuroheadset

The Emotiv EPOC+ 14 channels is an EEG neuroheadset that can produce measurable electric potentials to assess brain activities [35]. It is equipped with 14 saline sensors (electrodes) which are put on the scalp of the brain according to the international 10 to 20 system. In the 10 to 20 system, the real distance among two adjacent sensors can be either 10% or 20% [35]. The electrodes are situated in *F*_3_, *F*_4_, *FC*_5_, *FC*_6_, *F*_7_, *F*_8_, *AF*_3_, *AF*_4_, *T*_7_, *T*_8_, *O*_1_, *O*_2_, *P*_7_, *and P*_8_ locations and two reference electrodes-Driven Right Leg (DRL) and Common Mode sense (CMS) are located at *P*_3_ and *P*_4_ locations [35, 14, 17]. The electrodes distribution of Emotiv EPOC+ neuroheadset, employing the 10-20 system is shown in Fig. 1.

**Figure 1:**
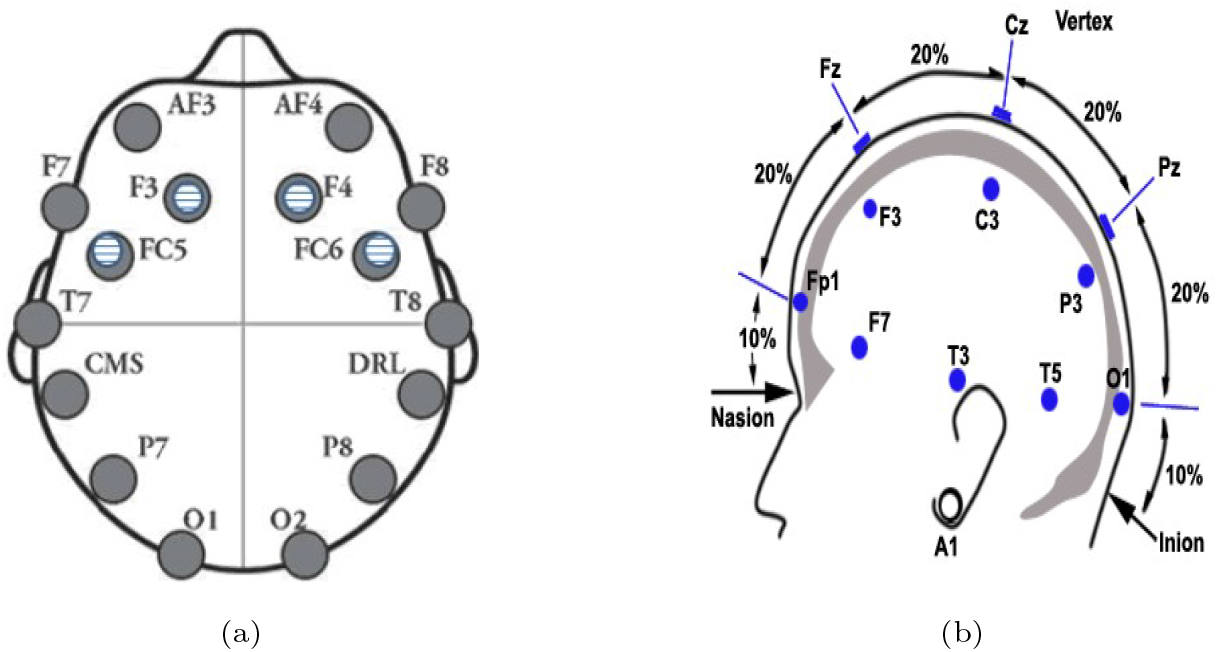
(a) Electrodes distribution of Emotiv EPOC+ neuroheadset (b) following the international 10-20 system. Source: Emotiv systems.

### 3.4. Signal Acquisition, Pre-processing, and Feature Extraction

In this study, twenty healthy subjects (age 18 ± 8) had participated with no prior neurological disorders. They were informed about the experimental procedures and gave informed consent before the experiment. Firstly, we developed an application program via Java-based technology inheriting Emotiv SDK and utilizing SDK API to acquire raw EEG brain signal from Emotiv neuroheadset. We get the European Data Format (EDF) formatted data from the SDK API. After that, we convert this file to CSV format via our program where we get the different frequency band values. This program along with CluSem and CluGame is also available at https://github.com/MdOchiuddinMiah/MI-BCI_ML/tree/master/Brain-Game/eeg_data_recording

As we mentioned earlier in Section 3.1, the primary motor cortex produces neural impulses that control the execution of movements. To collect this signal for our MI hand movement experiment, we had laid the headset based on the international 10-20 system. According to the electrode distribution in Fig. 1 and user manual of Emotiv which is available at https://emotiv.gitbook.io/epoc-user-manual, neuroheadset electrodes: *F*_3_, *FC*_5_, *FC*_6_, and *F*_4_ located on the frontal lobe which is familiar as the primary motor cortex of the brain. Furthermore, we had assessed the MI hand movement brain signal of 12 sensors without two reference sensors: *P*_3_ and *P*_4_. From the experiments and our prior works [14, 17], we had elected these sensors which are fitted in the primary motor cortex area of the brain. Additionally, these four electrodes are not placed in the exact same position where *C*_3_, *C*_4_, and *C*_*z*_ channels—these channels are commonly used for MI-based BCI to record important characteristics of MI—are located [3].

Also, this neuroheadset amalgamates a signal amplifier, 0.16Hz C-R high-pass filter, 85Hz analog low-pass filter, and a digital notch filter between 50Hz and 60Hz [35, 36]. This notch filter assists to neutralize the high-frequency noise in the signal. A sequential sampling at 128 samples per second is employed to convert an analog signal to digital by Delta-Sigma ADCs filtering [35, 37]. To acquire brain signal, we had connected the headset with the developed program computer through a wireless connection to the USB dongle and collected MI-EEG data for binary and ternary classes—the class values are steady, left, and right–hands. For each trial, visual cues on a computer screen with arrows pointing left and right were displayed for 30 seconds. These periods were interleaved for 2 seconds of fixations cross and 2 seconds of rest. Fig. 2 shows the 30 seconds of the visual cue of the training set and 15 seconds of the visual cue of the testing set for each trial. We collected the data from twenty healthy subjects. From each subject, we had taken two trials and one trial of each class to engender training and testing points, respectively. More details about our recorded training and testing datasets are also available at https://github.com/MdOchiuddinMiah/MI-BCI_ML/tree/master/Datasets.

**Figure 2:**
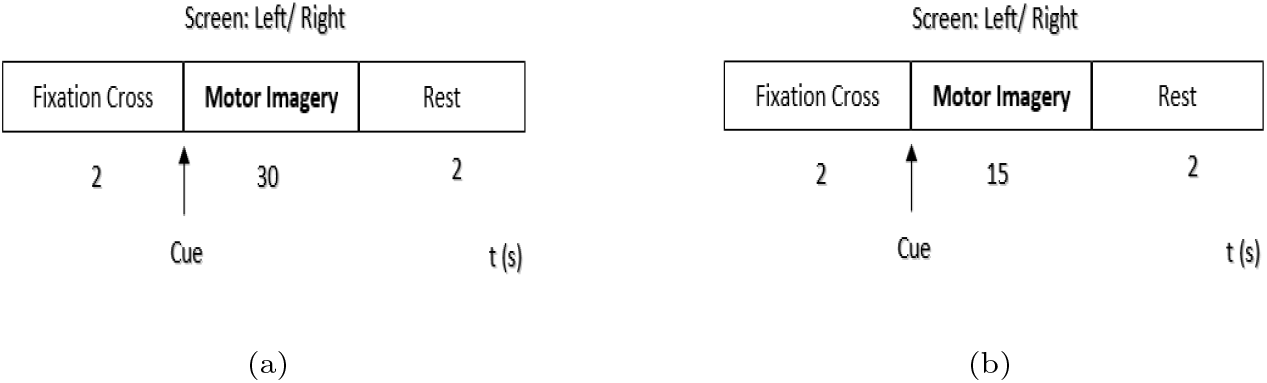
Experimental paradigm. One trial of the MI experiment. (a) Training points for 30 seconds (b) Testing points for 15 seconds.

To compute the band power of different frequency bands, we first calculate the Power Spectral Density (PSD) between band 4 to 45 Hz. PSD is a positive real function that is associated with a stationary stochastic process to measure the power strength at each frequency band [37, 38]. The fast Fourier Transform (FFT) method is directly used to compute PSD [14, 39]. By using a 2 seconds window size and 0.5 seconds step size, the average BP is calculated from the PSD of Theta (*θ*), Alpha (*α*), Low beta (*β*), High beta (*β*), and Gamma (*γ*) [14, 17, 27, 40]. Fig. 3 reveals the procedures how we acquire raw EEG data and prepare it to train the classification models through pre-processing and feature extraction techniques. Also, we extract features from the frequency bands, we compute the average band power (BP) of Theta (*θ*), Alpha (*α*), Low beta (*β*), High beta (*β*), and Gamma (*γ*) EEG neural frequency rhythms via the application program. EEG brainwaves, ranges, and their association with different activities in the brain are described in Section 3.5. Also, data pre-processing and band power calculation codes which we employed are available at https://github.com/MdOchiuddinMiah/MI-BCI_ML/tree/master/Pre-Processing.

**Figure 3:**
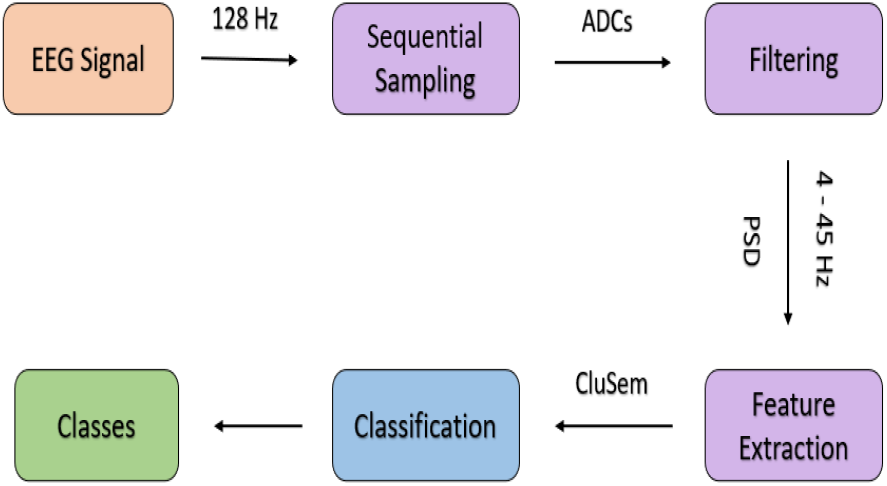
Sampling raw EEG data 128 samples per second, filtering with Delta-Sigma ADCs filtering, and calculate the PSD between band 4 to 45 Hz.

### 3.5. EEG Data Descriptions

The EEG measures different neural frequency rhythms that are associated with different regions, pathologies, or brain states. Neural frequency is assessed by calculating the number of waves repeats within a second [13, 41]. Table 1 illustrates the EEG brainwaves, their ranges, and their association with different activities in the brain [13, 3]. Most of the brain oscillations are connected with motor and sensory actions and associated with different brain functions [42, 34].

**Table 1:** EEG brainwaves, ranges, and their association with different activities in the brain.

| Rhythms | Band (Hz) | Different band activities inside the brain |
| --- | --- | --- |
| Theta ($\theta$) | 4 to 8 | Sleeping, drowsiness, meditation, relaxed |
| Alpha ($\alpha$) | 8 to 12 | Calmness, learning, body integration, relaxed |
| Low_beta ($\beta$) | 12 to 18 | Active concentration, arousal, conscious thought |
| High_beta ($\beta$) | 18 to 25 | Anxiety, task-oriented, logical thinking |
| Gamma ($\gamma$) | 25 to 45 | Cognitive functioning and high processing tasks |

We have divided the recorded MI-EEG dataset into binary-class and ternary-class sets based on the class values. We also collected an EEG eye state dataset from the UCI Machine Learning repository at https://archive.ics.uci.edu/ml/datasets/EEG+Eye+State to investigate the generality of our proposed clustering-based ensemble method. This EEG dataset is collected via an Emotiv EPOC+ neuroheadset from the EEG measurement of two different eye states [37]. The comprehensive information of binary-class, ternary-class, and EEG eye state datasets are presented in Table 2.

**Table 2:** Hand movement MI-EEG and EEG eye state datasets distribution.

| Datasets | Attributes<br>Number | Attribute<br>Types | Training<br>Data Points | Testing<br>Data Points | Classes |
| --- | --- | --- | --- | --- | --- |
| Binary-class<br>MI-EEG | 5 | Real &<br>Nominal | 22,800 | 5,400 | 2 |
| Ternary-class<br>MI-EEG | 5 | Real &<br>Nominal | 34,200 | 8,100 | 3 |
| EEG eye state | 15 | Real &<br>Nominal | 14,980 | 4,494 | 2 |

### 3.6. Visualization Brain Activities

The motor imagery hand movement deviations are found to be very high in the recorded MI-EEG responses as demonstrated in Fig. 4. We used the mean value of the five band power (BP) of different frequency bands to visualize the three activities. In this figure, we can see the average band power (BP) is higher for the activity of hand movements than the steady-state. It represents that the amplitude deviation between the movements and the steady is large. However, classifying this pattern for left and right-hand movements is a challenging task. As mentioned in Section 3.1, the primary motor cortex produces neural impulses to control the execution of movements left and right-hands. These movements are produced from the related area and generate similar patterns that introduce difficulty to classify, effectively.

**Figure 4:**
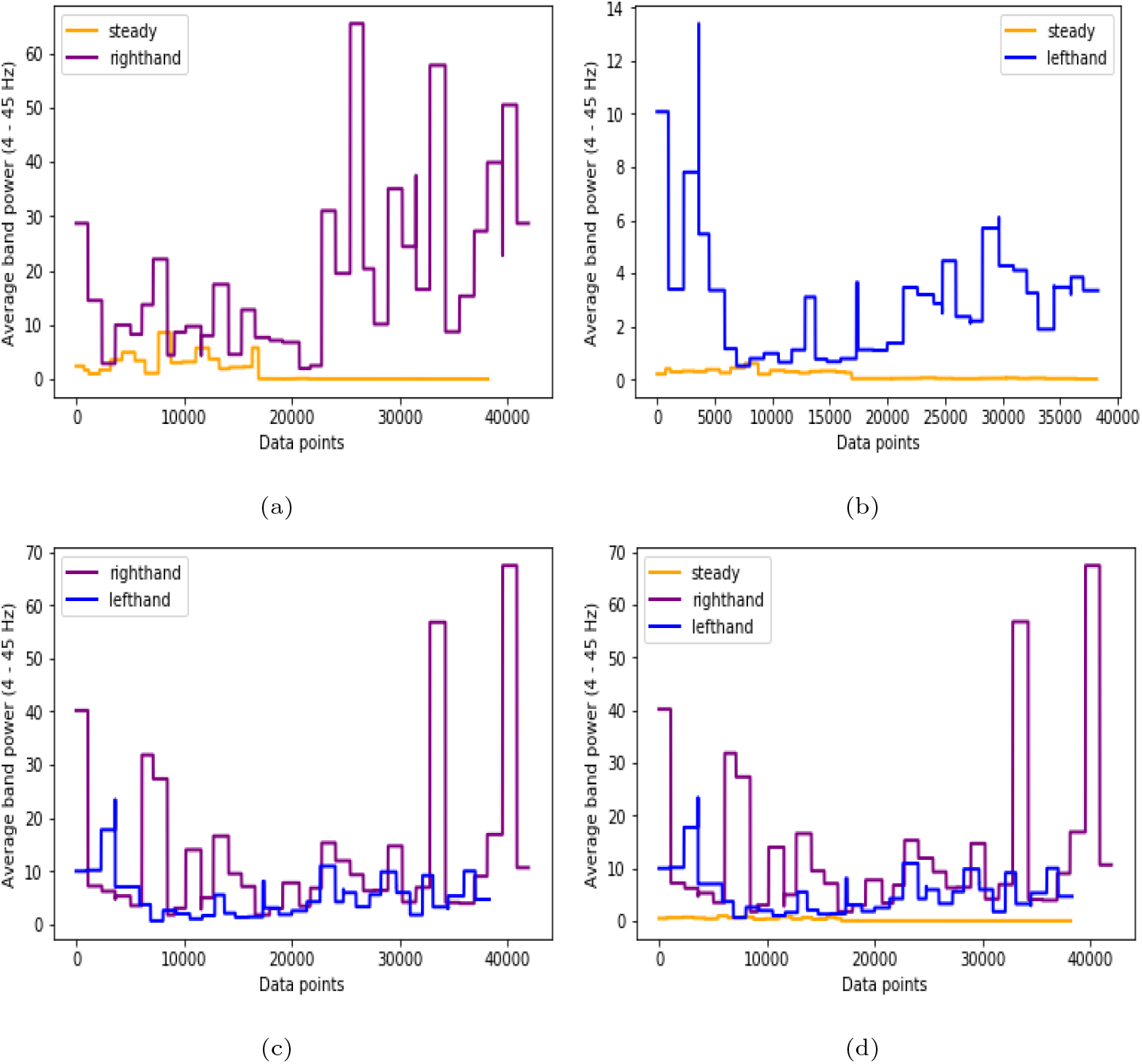
Dissimilar events inside the brain: (a) steady and right-hand movement; (b) steady and left-hand movement; (c) right and left-hand movement; (d) steady, right, and left-hand movement.

## 4. Classification

Classification task consists of training and testing steps [14]. In the training step, the labeled data points are used to train a classifier and generate a model. In the testing step, the unseen and unused data points are classified using the trained model to investigate its performance [43]. In this section, *N* is the number of instances in the dataset *D*, and *D* = {*x*_1_, …, *x*_*N*_} where each data point *x*_*n*_ ∈ *D* contains *F* features (*A*_*f*_, *f* = 1, …, *F*). In addition, 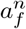 of data point *x*_*n*_ and each one belongs to a given class of *C* among {*c*_1_, …, *c*_*l*_, …, *c*_*M*_}. Here, we will briefly describe our proposed clustering-based ensemble model with four single classifiers and three ensemble learning methods.

### 4.1. Artificial Neural Network (ANN)

An artificial neural network (ANN) is a brain-inspired classification system based on the arrangement of biological neural networks [44]. It consists of an input layer, an output layer, and sometimes single or multiple hidden layers that exist to find complex patterns. The network contains several connections. Each connection computes the output of a neuron and then used by another neuron as input [30, 45]. Weights are allocated to each connection to reveal relative importance. Initially, all the weights are assigned randomly. After that, each neuron learns from the set of training instances *D*, computes the error according to the desired output of *D*, and adjusts the weights based on errors, accordingly. The backpropagation algorithm is employed updating weights to ameliorate the network until achieving the pure output weight [44, 46]. In this study, we use a Multi-Layered Perceptron (MLP) which is a feedforward artificial neural network (ANN) [44, 47]. Our employed MLP consists of one hidden layer and 100 nodes in this layer. We used one MLP hidden layer with 100 nodes, a learning rate 0.001, and 200 as the maximum number of iterations to enhance the classification performance.

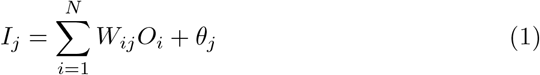

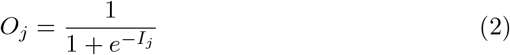

To compute neuron input *I*, it uses bias weight *θ*, connected all neurons weight *W* and output *O* [44]. After that, neuron input *I* manipulates to calculate neuron output *O* employing different activation functions. In this study, neuron input *I* and output *O* are calculated by employing non-linear sigmoid function which are illustrated in Eq. 1 and 2 [30, 48, 49].

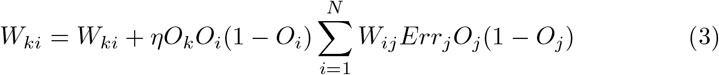

In this model, Eq. 3 is employed iteratively to adjust all the weights of the network. Here, the learning rate, *η* represents the proportion of corrective steps to adjust the errors of the model in each iteration [44, 30].

### 4.2. Support Vector Machine (SVM)

A Support Vector Machine (SVM) is a supervised classifier that explores training instances and discovers hyperplanes, or support vectors to maximize the margin among the classes [50]. In 2-dimensional space, the support vectors split a plane into two chunks through a line where each cluster denotes individual classes [19, 45]. A set of training data points, *D*= {*x*_1_, …, *x*_*N*_} having *N* number of data points with class values *C*= {1, −1} is employed to train the classifier. In this model, we select two parallel support vectors with supreme probable distance to separate classes. The maximum distance between two hyperplanes is called the margin [19]. A linear kernel is used with maximum iterations number to execute the algorithm.

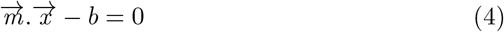

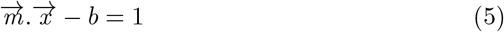

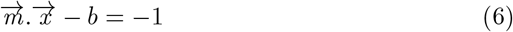

Any hyperplane can be described as the Eq. 4 where 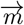 is a normal vector and parameter 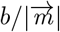 regulates the offset of the support vector from the origin [50]. Equations 5 and 6 are employed when the data points *x*_*n*_ are on or above the hyperplane and on or below the hyperplane respectively [22].

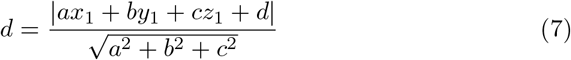

The distance, *d* among these two support vectors is 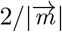. Hence, we have to minimize the 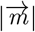 to maximize the *d*. The distance is calculated by manipulating the distance among a point and plane equation that are presented in Eq. 7 [50, 19].

### 4.3. Naïve Bayes Classifier (NB)

Naïve Bayes classifier is a probabilistic machine learning classifier based on Bayes’ theorem. It produces probability scanning the training instances only once and can handle the missing attribute values easily by omitting their probabilities [51, 45]. It takes a dataset, *D* with *F* number of attributes as input to build the classifier model. For testing a new instance, *x*_*new*_, the classifier will calculate the posterior probability and assign a class label with the highest probability.

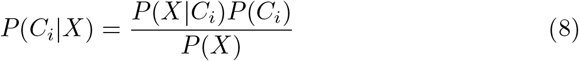

In Eq. 8, Bayes’ theorem states mathematically as *P* (*X*) is constant for all classes [51, 52]. Here, *P* (*X*|*C*_*i*_) and *P* (*C*_*i*_) represents conditional and priori class probabilities, respectively. For instance, NB classifier predicts a new data point, *X* as class *C*_*i*_, if *P*(*X*|*C*_*i*_)*P*(*C*_*i*_) is greater than *P*(*X*|*C*_*j*_)*P* (*C*_*j*_) in a binary classification task [43].

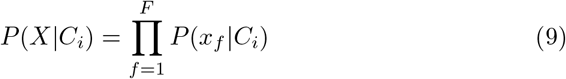

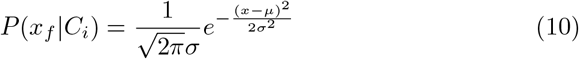

For data point *X, x*_*f*_ denotes the value of feature *A*_*f*_. The probability *P*(*x*_*f*_ |*C*_*i*_) can be simply assessed from the training instances, if feature, *A*_*f*_ is categorical-valued. Otherwise, if feature *A*_*f*_ has a continuous-value, then *A*_*f*_ is estimated through a Gaussian distribution with standard deviation *σ* and mean value *µ* shown in Eq. 10 [51].

### 4.4. Classification and Regression Tree (CART)

The CART uses *Gini Index* that produces binary classification tree to make decisions [14]. Firstly, it assesses the adulteration of dataset, *D* where probability *P*_*n*_ is estimated through |*c*_*l*_, *D*|*/*|*D*| presented in Eq. 11 [51]. Here, the sum is calculated over *C* classes and each instance, *x*_*n*_ ∈ *D* belongs to a class *c*_*l*_.

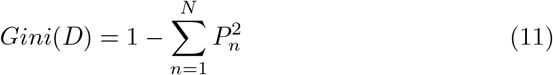

It splits the dataset, *D* considering binary split and the weighted sum of the adulteration of every resulting sub-data. For instance, the dataset, *D* splits into *D*_1_ and *D*_2_ considering the *Gini Index* of *D* which is calculated in Eq. 12 [51].

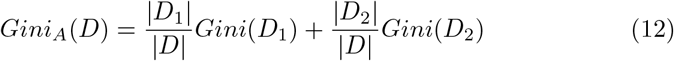

For each attribute, *A*_*f*_ considers each probable sub-data and is used as the splitting attribute which ameliorates the reduction impurity, as shown in Eq. 13 [51].

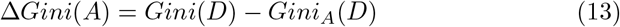

According to the *Gini Index* impurity, it will split the dataset and create leaf nodes until all the splitting data belongs to their corresponding classes. Algorithm 1 outlines the Classification and Regression Tree Algorithm [51]. To achieve the best result, the maximum depth of the tree and the best splitting technique used at each node.

### 4.5. Random Forest (RF)

Random forest is an ensemble approach to classify a high volume of data with superior accuracy [43]. Initially, it splits a set of instances, *D* into several subsets (*D*_1_, *D*_2_, …, *D*_*n*_) depending on the number of features *F*. Then, it constructs multiple models by using a learning scheme based on the number of subsets and classifies samples in each subset [43, 20]. The learning scheme that is used to build RF is presented in Algorithm 1. In RF, there is an association between the accuracy and the number of generated models. In other words, the number of models is directly proportional to accuracy. In machine learning, overfitting is one of the crucial problems and it may abate the classifier accuracy [43]. RF classifier is among the classifiers that are least prone to this problem. As it considers the vote of every model, it is less likely to overfit the model and usually gain greater accuracy than a single classifier [53, 43]. The number of trees in the forest, 100 is used with maximum depth to achieve better accuracy.

#### Algorithm 1 CART Algorithm

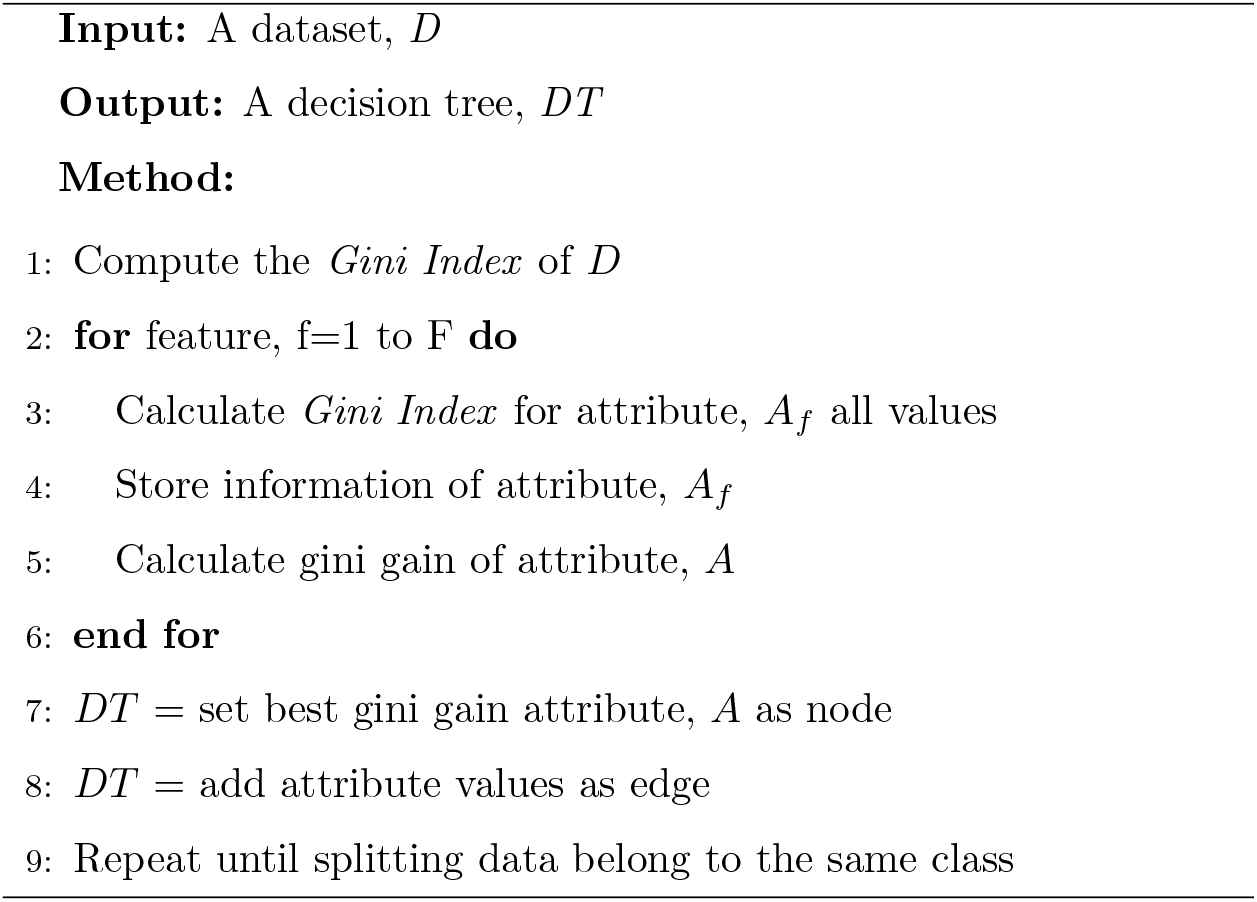

### 4.6. Bagging

Bagging is also called Bootstrap Aggregation, which is an ensemble approach employed in statistical classification and regression to boost the performance of ML algorithms [1]. It combines the prediction of different equal-weighted models and classifies a new instance using different voting technique[20]. The bagging technique requires a set of instances *D* of size *N* and several iterations, *I* to build the classifier model as its parameters. It generates new subsets, ({*D*_1_, *D*_2_, …, *D*_*n*_}) by sampling the original dataset, *D* with replacement until the iterations number. Then, it uses each sub-set and learning scheme illustrated in Algorithm 1 to derive classifier models. To classify a new instance *x*_*new*_, it combines the output of each model and uses majority voting to combine the prediction results [1]. In the ensemble, the number of base estimators, 10 used with the CART algorithm.

### 4.7. AdaBoost

Adaptive Boosting (AdaBoost) is a popular ML meta-algorithm, which combines a series of classifiers weighted votes to classify instances [1, 52]. It builds a strong classifier by combining a set of weak classifiers [54]. Initially, it assigns an equal weight, 1*/d* to each training data point, *x*_*i*_ ∈ *D*. Then, it produces a group of subsets, ({*D*_1_, *D*_2_, …, *D*_*n*_}) by sampling *D* with replacement based on instance weight until the iterations number, *I*. Each generated dataset, *D*_*i*_ derives a model, *M*_*i*_, and computes the error of the model by adding the weights of all instances in *D*_*i*_. Eq. 14 shows the error calculation function of a model where *err*(*x*_*j*_) will be 1 when *x*_*j*_ is misclassified, and 0 otherwise [55].

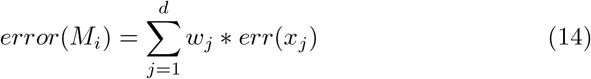

If the error of a model exceeds 0.5, we will regenerate *D*_*i*_ and derive a new *M*_*i*_. We will update the weight of an instance as if the weight of classified instances is abated and misclassified instances are enlarged [51, 52, 53]. To classify a new instance, it combines the votes and weights of each classifier. The base estimator CART is used with maximum depth and 100 estimators to execute AdaBoost for best classification results.

### 4.8. Proposed Ensemble Method

An Ensemble method is a technique to construct a model by incorporating several classifiers to achieve better performance. The proposed clustering-based ensemble technique CluSem that is used to classify motor imagery hand movement will discuss in this section. Initially, the proposed method takes the training EEG brain data *D*= {*x*_1_, …, *x*_*N*_} from *NE* neurons/ electrodes. It then divides the dataset, *D* into *NE* clusters based on the label of the electrodes. All clustered datasets {*D*_1_, *D*_2_, …, *D*_*NE*_} represent individual electrodes as we divide dataset, *D* based on it. Again, each of these clustered datasets divides into *C* clusters based on the attributes of each subset. Finally, we get *NE* datasets after the first clustering and the end of the second clustering, we get *C* subsets {*D*_1_, *D*_2_, …, *D*_*C*_} for each subsets {*D*_1_, *D*_2_, …, *D*_*NE*_}.

After generating the subsets, we construct *C* number of decision trees *DT*_1_, *DT*_2_, …, *DT*_*C*_ for *m*^*th*^ cluster using decision tree induction algorithm (CART). The CART algorithm is explained comprehensively in section 4.4. We compute the error rate of *DT*_1_, *DT*_2_, …, *DT*_*C*_ on dataset, *D*_*n*_ and find the decision tree, *min*_*DT*_ ∈ *DT*_*m*_ with the minimum error rate *min*_*error*_. Then, this tree *DT*_*m*_ is considered to construct the final ensemble of trees *DT*\* and models *M*\*. In the last step of this CluSem method, each MI-EEG data point, *x*_*new*_ classifies with *M*\* based on the position of the electrodes and deliberate the majority vote as prediction among the classification results of *M*_*n*_ ∈ *M*\*. Here, the data points are coming from different electrodes and select the model dynamically based on the position of the electrodes.

Before incorporating the CART as a learning scheme in CluSem, we had examined the performance and time complexity by using other popular methods in CluSem. The methods which were tested besides CART are ANN, SVM, Naïve Bayes, RF, Bagging, and AdaBoost. The classification performance and execution time of this experiment are illustrated in Table 3. The primary reason to do this experiment is to find the best parameter for our proposed method which provides good results with less computational time. As we employ this proposed method in our application to classify real-time data, we need a method that can be able to give good results with less execution time. To get a better user experience, we have to make our application fast and accurate. For this, we used CART in our proposed method which is simpler than other algorithms—they are evaluated in Table 3. The process of classifying motor imagery tasks is illustrated in Fig.5. Also, the proposed clustering-based ensemble method CluSem is summarized in Algorithm 2.

**Table 3:**
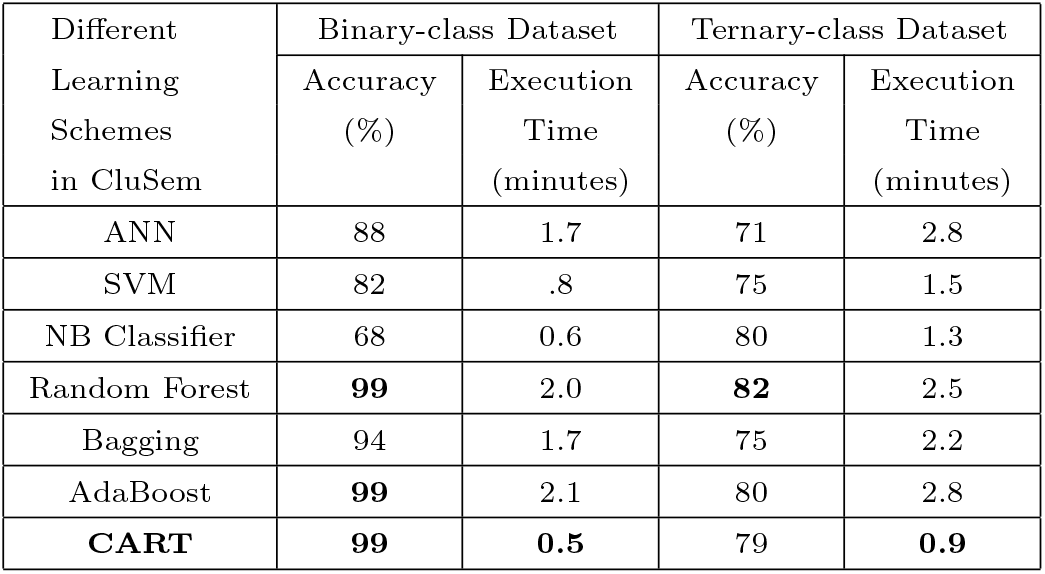
The results reported using different classifiers as a learning scheme of the CluSem model using binary-class and ternary-class datasets.

**Figure 5:**
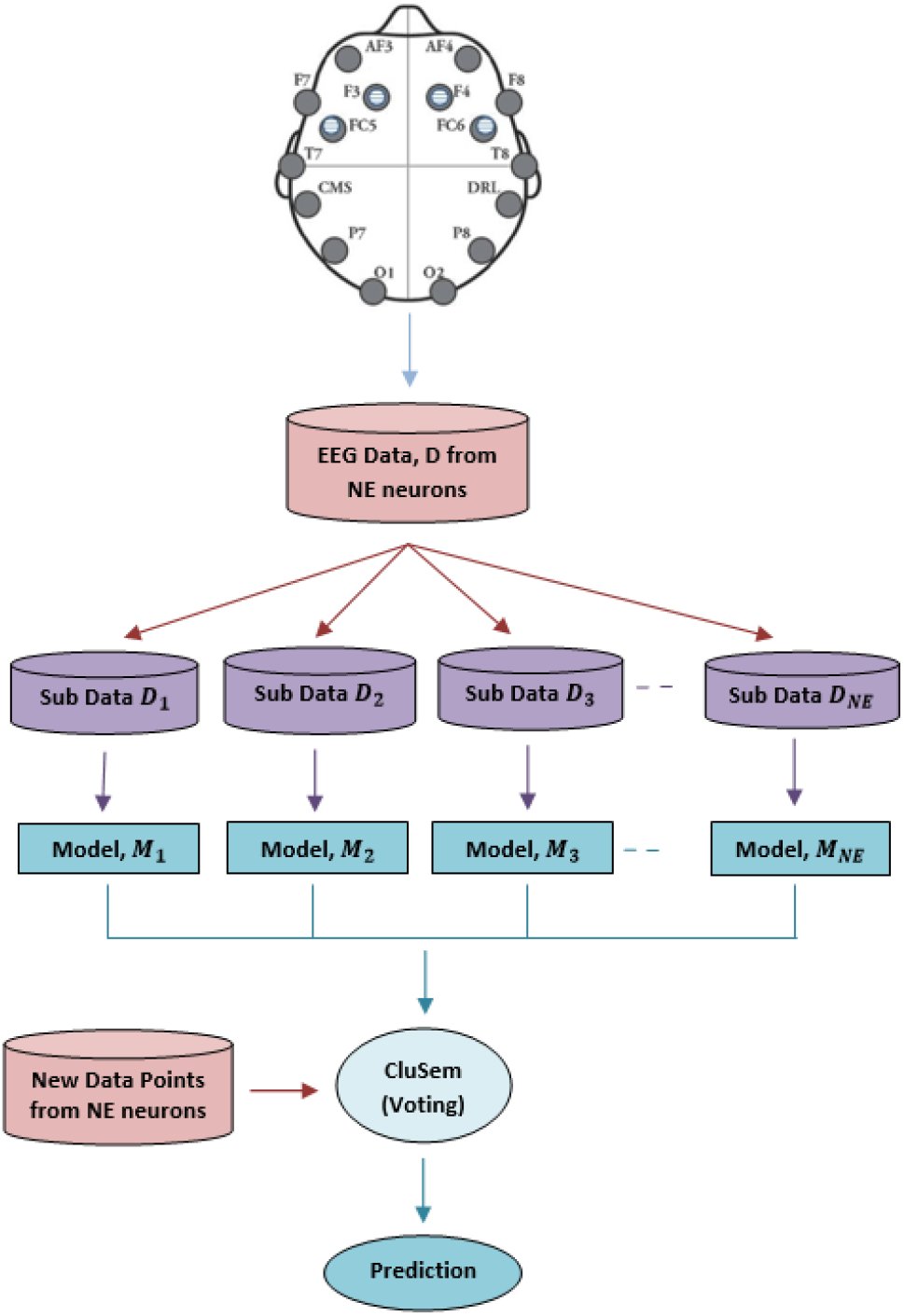
The process of CluSem method to classify real-time motor imagery EEG data.

In BCI, real-time MI related EEG signal classification is a challenging task. Most often, the signals are biased with artifacts and noise due to the low conductivity of the electrodes with the scalp [1, 24]. Different areas of the brain are responsible for the individual task and each electrode on the scalp provides different signals [20]. Enhancing the performance of EEG signal classification in real-time is a demanding task because of the high dimensionality of the input data and dynamic behavior of the electrodes [54].

#### Algorithm 2 Proposed Clustering-based Ensemble Method

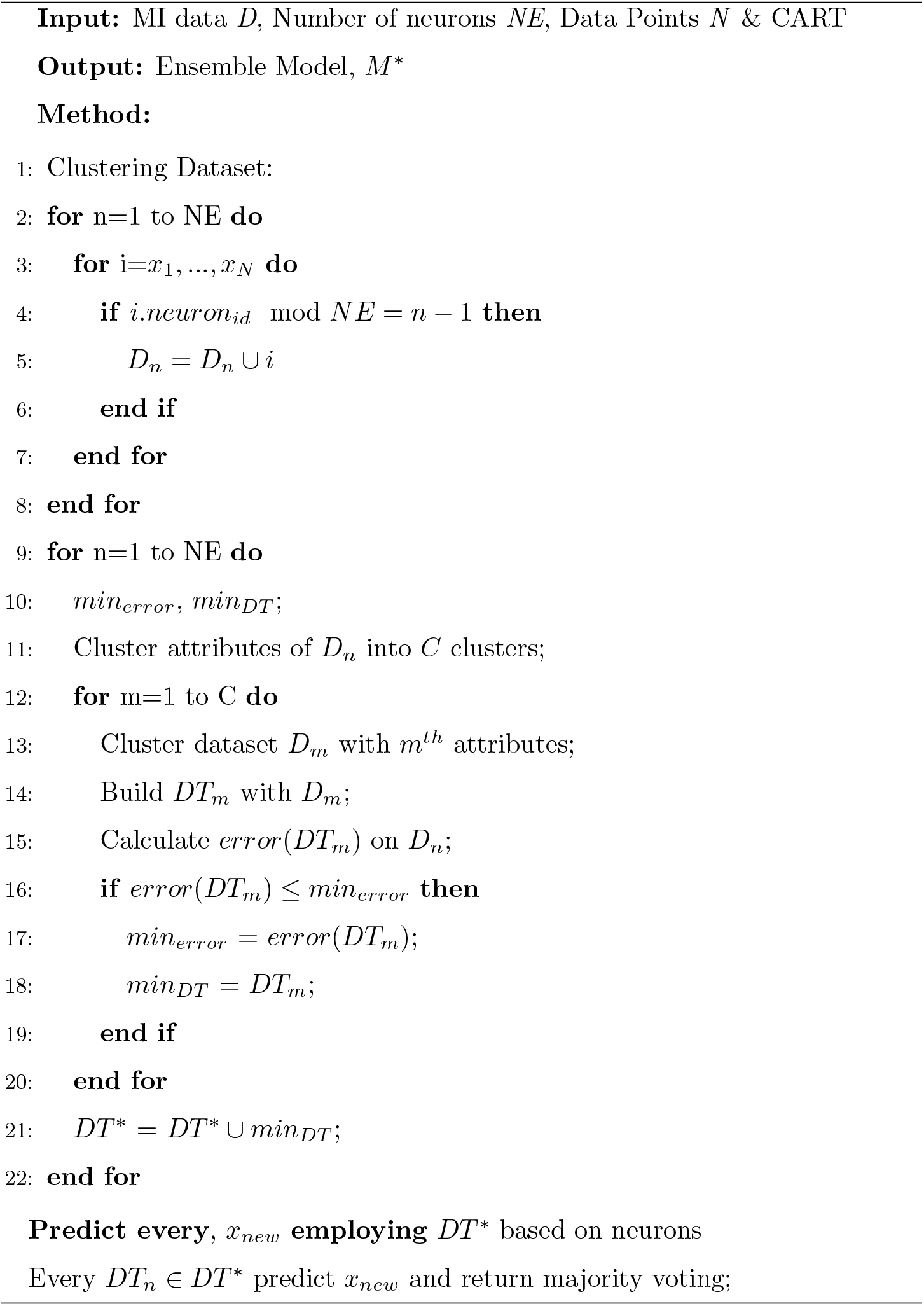

It is a challenging task to build a good model employing single classifiers like ANN, SVM, naïve Bayes, or Decision Tree for high-dimensional input data [20, 21]. On the other hand, ensemble learning methodologies are employing widely to tackle this issue and demonstrated promising results. However, the existing ensemble methods generate subsets by sampling the original dataset with replacement technique [1, 24]. By applying this technique, the same instance can be repeated several times. The idea behind the clustering approach before the classification that is proposed in this study is to overcome this problem and to classify MI related EEG signals in real-time. It clusters the dataset based on the position of the electrodes, hence each cluster represents dissimilar information. It also selects the model dynamically based on the electrode locations to classify real-time EEG data. In this study, we have used several classifiers that are widely used in this field to investigate the effectiveness of our proposed method [1, 24, 28]. To do this, we used some single classifiers: ANN, SVM, NB Classifier, CART, and popular ensemble classifiers: RF, Bagging, AdaBoost which are employed in recent days [14, 1, 2, 28, 7, 56].

## 5. Experiments

In this section, we present the experimental environment, results of the proposed clustering-based ensemble method, and our developed brain game that is controlled by real-time motor imagery hand movements data.

### 5.1. Experimental Setup

In this study, the experiments were conducted via a device with an Intel Core-i5 (2.60 GHz) processor, and 8 GB of RAM. We implement the proposed method in Python programming language (version 3.7) and used the scikit-learn (version 0.21.2) as a machine learning library. We compare the performance of the proposed method with several popular machine learning algorithms using classification accuracy, precision, recall, and F-score.

The accuracy is measured by Eq. 15 where *assess*(*x*_*i*_) = 1 when *x*_*i*_ is correctly classified or *assess*(*x*_*i*_) = 0 when *x*_*i*_ is misclassified [53]. The precision, recall, and F-score are calculated using Eq. 16 to 18, respectively [51]. We also represent decision boundaries using the area under the ROC curve and AUC score. In the ROC curve, true positive rate (TPR) is plotted against false positive rate (FPR). The calculations are presented in Eq. 19 and 20. In au-ROC curve, the lowest threshold is considered through the *y* = *x* line where correctly classified data points are represented by 1 and misclassified instances are represented as 0 [43].

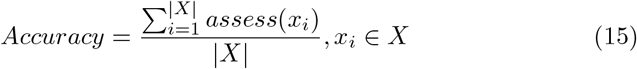

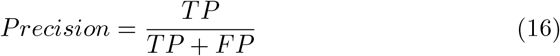

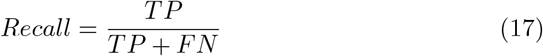

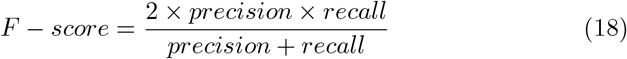

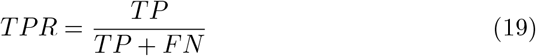

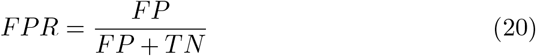

Here, TP, TN, FP, and FN indicate the number of positive samples correctly classified, negative samples correctly classified, negative samples incorrectly classified, and positive samples incorrectly classified respectively [53, 14].

### 5.2. Results

In the first step, we evaluate the performances of CluSem with different learning schemes—they are some popular machine learning classifiers—used in this experiment. We mainly scrutinize the CluSem performance and the execution time needed to test the model with different parameters. The result of this comparison is presented in Table 3 which is examined by two datasets—binary-class and ternary-class datasets—with algorithms: ANN, SVM, NB Classifier, RF, Bagging, AdaBoost, and CART. Among these algorithms, CART needs less computational time and provides better accuracy in both datasets. RF, Bagging, and AdaBoost also provide good results but failed to execute with less execution time.

In the second step, we evaluate the performances of CluSem against existing methods that are widely used for this task namely, ANN, SVM, Naïve Bayes, CART, RF, Bagging, and AdaBoost Classifiers on training sets of motor imagery EEG datasets. The results for this comparison is presented in Table 4. The results in this table demonstrate that the proposed clustering-based ensemble algorithm significantly outperforms the existing classifiers on a binary-class motor imagery EEG dataset. The proposed method performed superior and reaches 99.0% in terms of accuracy for the binary-class dataset. In this dataset, single classifiers are failed to achieve more than 85.0% accuracy where existing ensemble approaches can achieve up to 91.0% accuracy. As shown in this table, CluSem can achieve over 8.0% better performance compared to other ensemble classifiers used for this task. The decision boundaries of the proposed method compared to employed classifiers for this dataset is shown in Fig. 6.

**Figure 6:**
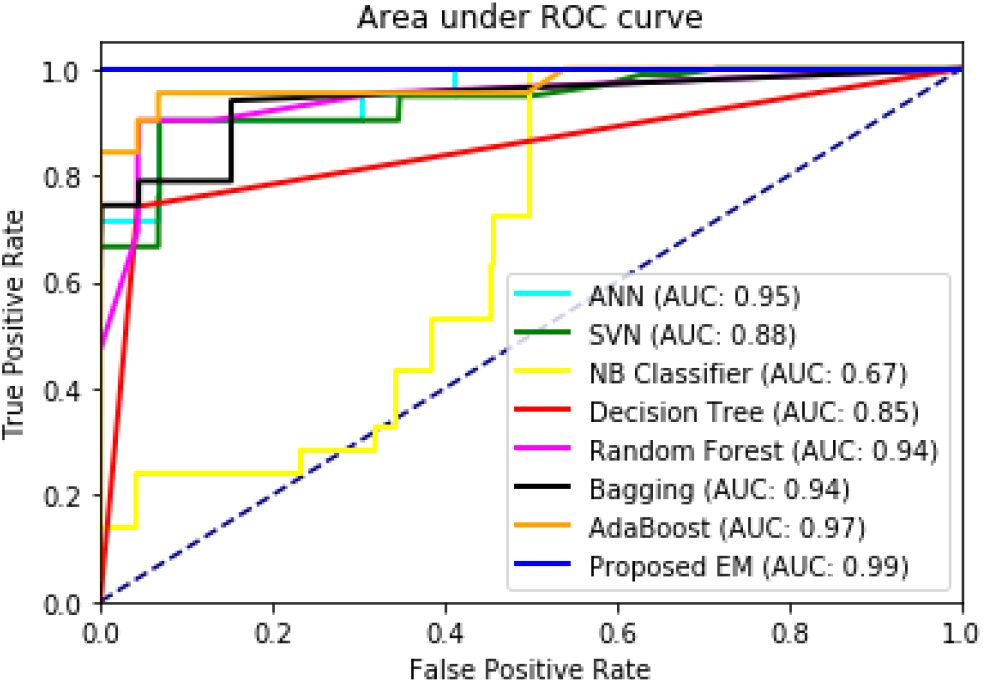
ROC and AUC analysis of CluSem method compared to popular classifiers on binary-class motor imagery EEG dataset.

**Table 4:** The results reported using the CluSem model compare to popular classifiers on binary-class EEG dataset.

| Classifiers | Accuracy (%) | Recall | Precision | F-score |
| --- | --- | --- | --- | --- |
| ANN | 84 | 0.84 | 0.86 | 0.84 |
| SVM | 78 | 0.77 | 0.80 | 0.79 |
| NB Classifier | 60 | 0.60 | 0.68 | 0.62 |
| CART | 85 | 0.87 | 0.85 | 0.85 |
| Random Forest | 90 | 0.90 | 0.91 | 0.90 |
| Bagging | 82 | 0.82 | 0.82 | 0.82 |
| AdaBoost | 91 | 0.91 | 0.91 | 0.91 |
| **Proposed EM** | **99** | **0.99** | **0.98** | **0.97** |

Moreover, as shown in Table 5, the proposed clustering-based ensemble classifier also outperforms some popular machine learning algorithms on the ternary-class dataset. In the ternary-class dataset, left and right-hand movements classification is challenging to predict because both tasks are engendered from the motor cortex, and samples are closely associated. Similarly, our proposed method achieved better than the existing four single classifiers and three ensemble methods. As shown in this Table, CluSem achieves 79.0% prediction accuracy which is 17.0% better than those reported using other classifiers. Fig. 7 illustrates the decision boundaries of the classifiers via ROC analysis and AUC scores for the ternary-class dataset.

**Table 5:** The results reported using the CluSem model compare to popular classifiers on the ternary-class EEG dataset.

| Classifiers | Accuracy (%) | Recall | Precision | F-score |
| --- | --- | --- | --- | --- |
| ANN | 53 | 0.53 | 0.55 | 0.54 |
| SVM | 57 | 0.55 | 0.57 | 0.56 |
| NB Classifier | 60 | 0.60 | 0.64 | 0.59 |
| CART | 59 | 0.60 | 0.59 | 0.58 |
| Random Forest | 62 | 0.62 | 0.64 | 0.61 |
| Bagging | 57 | 0.57 | 0.59 | 0.57 |
| AdaBoost | 61 | 0.62 | 0.63 | 0.59 |
| **Proposed EM** | **79** | **0.79** | **0.80** | **0.79** |

**Figure 7:**
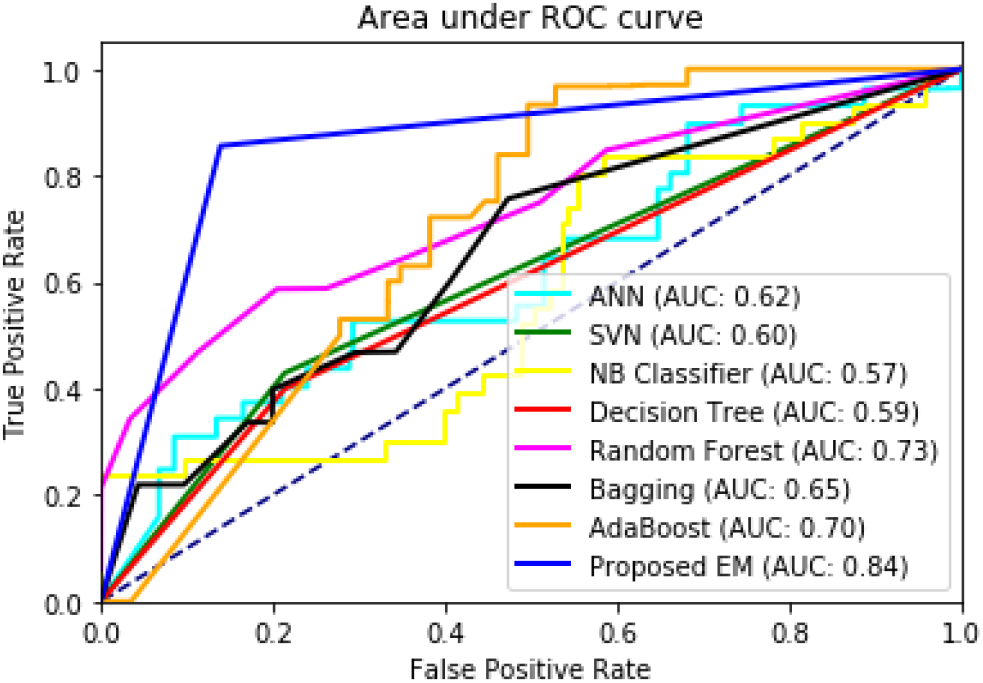
ROC and AUC analysis of CluSem method compared to popular classifiers on ternary-class motor imagery EEG dataset.

Besides, we used the EEG eye state dataset to experience our proposed clustering-based ensemble method. For this dataset, CluSem also achieves significant improvements over other classifiers. It achieves 90.0% accuracy on average while decision tree CART achieves 83.0% accuracy, and existing ensemble approaches achieve up to 89.0% accuracy. The comparison of accuracy, precision, recall, and F-score analysis using 10-fold cross-validation for the EEG eye state dataset is presented in Table 6. Fig. 8 illustrates the ROC analysis and AUC scores of the proposed method for the EEG eye state dataset.

**Table 6:** The results reported using the CluSem model compare to popular classifiers on EEG eye state dataset.

| Classifiers | Accuracy (%) | Recall | Precision | F-score |
| --- | --- | --- | --- | --- |
| ANN | 44 | 0.43 | 0.44 | 0.43 |
| SVM | 64 | 0.63 | 0.63 | 0.63 |
| NB Classifier | 54 | 0.55 | 0.55 | 0.42 |
| CART | 83 | 0.84 | 0.84 | 0.84 |
| Random Forest | 88 | 0.89 | 0.90 | 0.90 |
| Bagging | 89 | 0.90 | 0.90 | 0.90 |
| AdaBoost | 75 | 0.75 | 0.75 | 0.75 |
| **Proposed EM** | **90** | **0.90** | **0.91** | **0.90** |

**Figure 8:**
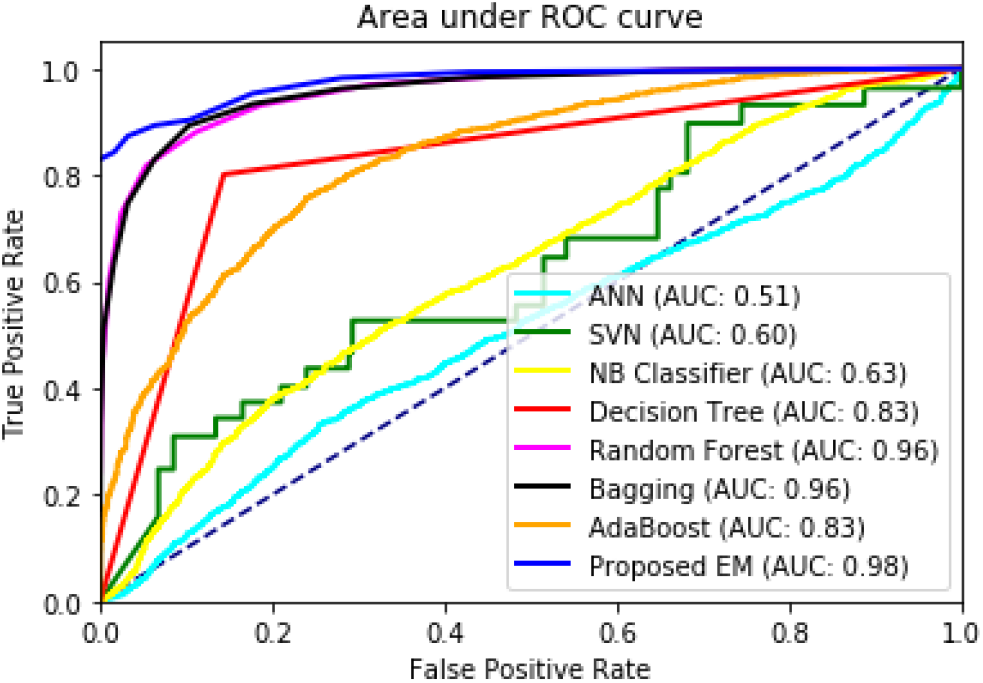
ROC and AUC analysis of CluSem method compared to popular classifiers on EEG eye state dataset.

Table 7 shows a comparison of our proposed work with the other state-of-the-art methods. To make a fair comparison, all the methods were implemented and evaluated using 10 × 10-fold cross-validation scheme on the EEG eye state dataset. It can be noted that the proposed method outperformed all the other state-of-the art methods in terms of accuracy, recall, precision and F-score. Our proposed method achieved an improvement in accuracy of 5.80% compared to the conventional common spatial pattern (CSP) approach and an improvement in accuracy of 7.28% compared to the recently proposed OPTICAL+ predictor. The SS-MEMDBF method also performed well as it has the 2^*nd*^ best performance. The OPTICAL+ predictor did not perform well, which might be due to low number of training samples as the OPTICAL+ predictor uses deep learning technique.

**Table 7:** 10 × 10-fold cross-validation results of the different methods on the EEG eye state dataset.

| Methods | Accuracy (%) | Recall | Precision | F-score |
| --- | --- | --- | --- | --- |
| CSP | 84.20 | 0.832 | 0.855 | 0.838 |
| DFBCSP [57] | 79.99 | 0.759 | 0.830 | 0.778 |
| I-DFBCSP [25] | 83.38 | 0.796 | 0.860 | 0.814 |
| OPTICAL [58] | 76.07 | 0.698 | 0.782 | 0.730 |
| OPTICAL+ [59] | 82.72 | 0.806 | 0.838 | 0.809 |
| SS-MEMDBF [60] | 89.36 | 0.891 | 0.889 | 0.884 |
| SFTOFSRC [40] | 73.03 | 0.769 | 0.708 | 0.730 |
| **Proposed EM** | **90.00** | **0.900** | **0.910** | **0.900** |

As shown in this Section, CluSem can outperform the most popular classification techniques that are used for this task. It can achieve high accuracy and AUC scores on average on three EEG datasets. CluSem delivers an enhancement between 10% and 20% accuracy compared to individual classifiers. It also achieves approximately 5% to 15% better prediction accuracy compared to existing ensemble approaches. Although the proposed clustering-based ensemble method outperforms other existing classifiers, there are limitations to be considered. It is possible that the input signal would be influenced by artifacts and noise due to the short conductivity of the neurons with the scalp. In such circumstances, CluSem or any other classifiers that are investigated in this study will not be able to correctly classify data points in real-time and the performance of the classifier is abated.

### 5.3. Developing Brain Game

We used our proposed clustering-based ensemble model (CluSem) and Java-Swing technology to develop our targeted application system called CluGame. We also used the Emotiv community SDK in our Java program to acquire live brain signals. This game takes the average band power of different frequency bands from *F*_3_, *FC*_5_, *FC*_6_, and *F*_4_ electrodes as input commands. Then these commands are classified by CluSem based on the position of the electrodes and generate actions as the prediction, accordingly. The input commands are coming from *F*_3_, *FC*_5_, *FC*_6_, *F*_4_ electrodes, and select the model dynamically based on the position of the electrodes.

In CluGame, we present several actions via animated balls according to the prediction accuracy of different classes. The real brain data classification through the developed model and prediction tabulation via animated balls in real-time is controlled via threads. The final prediction is taken from the dynamic number of instances votes to control the game more precisely. Here, we have developed two versions of this game: CluGame-2 and CluGame-3 for binary as well as ternary actions of motor imagery task without any necessity of conventional input devices, respectively. It also increased the classification accuracy of real-time EEG signals of motor imagery tasks.

A sample view of CluGame is shown in Fig. 9. The animated balls are presenting the movement and steady accordingly. In this figure, the balls are in two labels: movement and steady. When the game predicts the input as steady, the balls will be steady and label as steady. On the other hand, the animated balls will be in a movement state when it predicts the input as left- or right-hand movements. If there is any error or failure to predict or the headset is not working, CluGame will provide proper messages. In real life, this game can be used for rehabilitation as well as upgrading user well-being. The user can exercise her/ his concentration to recover from attention deficiency and boost her/ his attention via playing this brain game. It can also simply be used for gaming and entertainment. The source code of brain game is also available via open repository at https://github.com/MdOchiuddinMiah/MI-BCI_ML/tree/master/Brain-Game.

**Figure 9:**
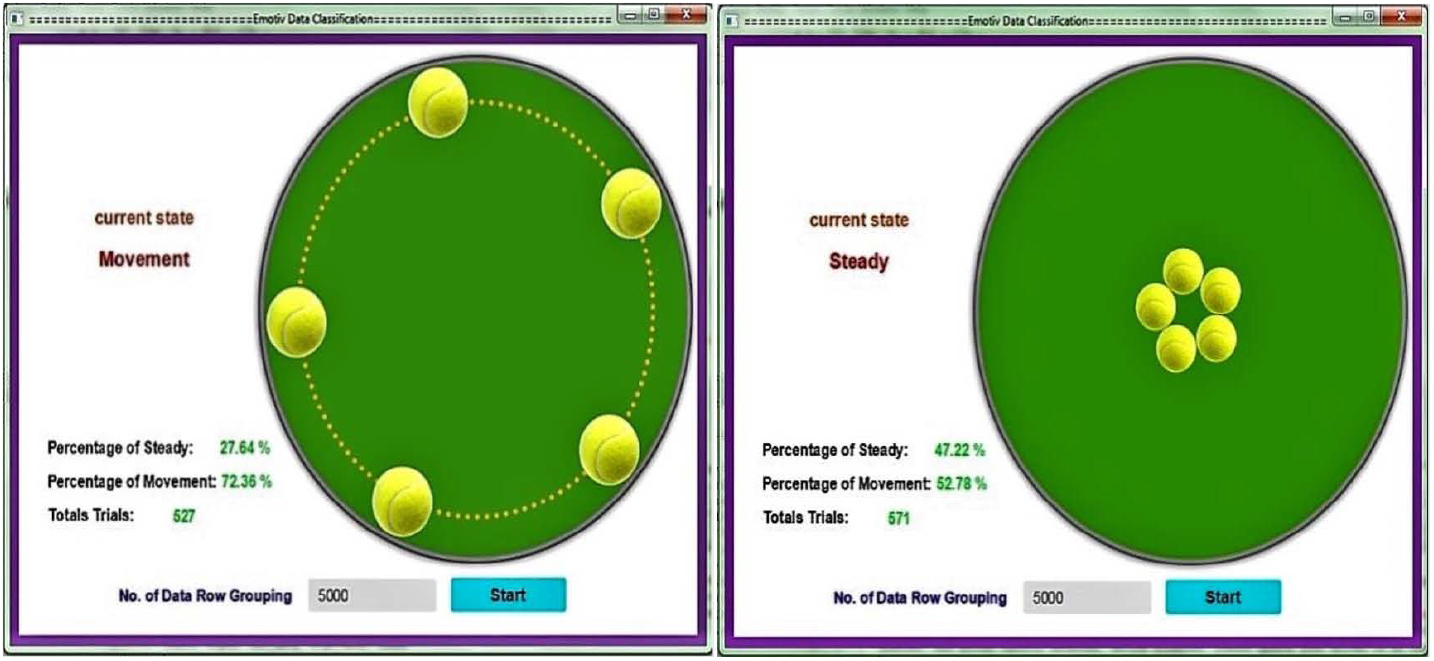
Classify real-time motor imagery tasks via developed CluGame, presenting movements and steady correspondingly.

## 6. Conclusion and Future Work

In this study, we proposed a new tool called CluSem to predict motor imagery tasks from multi-channel EEG Data. To build this model, we have used Emotiv EPOC+ EEG neuroheadset after analyzing several BMI devices like Emotiv EPOC+, Muse Headband, Aurora Dream, and MindWave. To obtain MI-EEG brain signals, we built an application program manipulating Emotiv SDK with Java technology.

We have investigated the performance of CluSem compared to the most popular classifiers used for this task such as ANN, SVM, Naïve Bayes, CART, RF, Bagging, and AdaBoost. Our results demonstrate that CluSem can significantly outperform previously used classifiers for this task. It is also able to enhance the prediction performance between 5% and 15% over the previously proposed ensemble classifiers to solve this problem.

Besides, we use CluSem to develop a game called CluGame that is capable of controlling the movements of the balls utilizing the real-time MI-EEG brain signals. It offers the user to enhance the quality of attention, which boosts productivity. It also assists people who are physically impaired or disabled and can potentially be used for human functionality enhancement. We designed the game to be a single-player game and to recognize three actions. In the future, more actions and more players can be added to make the game more advanced and challenging. The finding of this research can also be applied to manipulate and enhance the control as well as movements of robots. It also brings new potentials in the health and rehabilitation industry. CluSem, CluGame, our generated benchmark datasets, and all associated codes are publicly available at https://github.com/MdOchiuddinMiah/MI-BCI_ML.

## Ethical Approval

All applicable international, national, and/or institutional guidelines for the care and use of animals were followed. The ethical approval is taken from United International University which is available at https://github.com/MdOchiuddinMiah/MI-BCI_ML/blob/master/EthicalApproval.pdf

## Informed Consent

Informed consent was obtained from all individual participants included in the study.

